# *Bacteroides* is increased in an autism cohort and induces autism-relevant behavioral changes in mice in a sex-dependent manner

**DOI:** 10.1101/2023.10.05.560465

**Authors:** Julie Carmel, Nasreen Ghanayem, Rasha Mayouf, Natalia Saleev, Ipsita Chaterjee, Dmitriy Getselter, Evgeny Tiknonov, Sondra Turjeman, Mounia Shaalan, Saleh Khatib, Alla Kuzminsky, Neta Kevtani-Friedman, Tanya Kronos, Tali Bretler, Omry Koren, Evan Elliott

## Abstract

Autism Spectrum Disorder (ASD) is a neurodevelopmental condition which is defined by decreased social communication and the presence of repetitive or stereotypic behaviors. Recent evidence has suggested that the gut-brain axis may be important in neurodevelopment in general and may play a role in ASD in particular. Here, we present a study of the gut microbiome in 96 individuals diagnosed with ASD in Israel, compared to 42 neurotypical individuals. We determined differences in alpha and beta diversity in the microbiome of individuals with ASD and demonstrated that the phylum Bacteroidetes and genus *Bacteroides* were the most significantly over-represented in individuals with ASD. To understand the possible functional significance of these changes, we treated newborn mice with *Bacteroides fragilis* at birth. *B. fragilis*-treated male mice displayed social behavior dysfunction, increased repetitive behaviors and gene expression dysregulation in the prefrontal cortex, while female mice did not display behavioral deficits. These findings suggest that overabundance of *Bacteroides*, particularly in early life, may have functional consequences for individuals with ASD.

## Introduction

Autism Spectrum Disorders (ASD) are defined by dysregulation of social communication and the presence of repetitive or stereotypic behaviors^1^. Hypersensitivity or hyposensitivity to specific sensory inputs are commonplace in ASD as well. The etiology of ASD is defined by a strong genetic component. Several hundred autism-associated genes have been identified, but evidence has also highlighted a strong role for common variants in the development of ASD^2^ . Both twin studies and epidemiological studies suggest that there is a significant contribution of environmental factors as well. While it is not yet clear what these exact environmental factors are, several lines of evidence have implicated maternal immune activation during pregnancy, particularly the first trimester, as one potential environmental factor initiating ASD^3^.

The gut-brain axis, and particularly the gut-microbiota-brain axis, has been identified as a substantial influencer of mammalian behavior, including social behavior^4,5^. The gut microbiome may affect brain function through several pathways, including metabolites, regulation of the immune system, and direct influence on the central nervous system via the vagus nerve^6–8^. Preliminary studies in mouse models determined the possible influence of specific bacterial species, such as *Limosilactobacillus reuteri*^9^and *Bacteroides fragilis*^10^, on ASD-related behaviors. Parallel studies have also identified some dysregulation of the gut microbiome in individuals with ASD, though the exact bacterial changes observed vary considerably between the different studies^11^. One study determined that offspring of mice gavaged with gut microbiota from humans diagnosed with ASD displayed ASD-like behavior, thereby providing a possible causal link between human microbial dysregulation and behaviors associated with ASD^12^.

Most previous rodent studies that examined specific bacterial populations in ASD focused on uncovering dysbiosis in ASD rodent models during adulthood followed by microbiome treatment. However, there are few studies where specific changes in microbiome in individuals with ASD were modeled by treatment of mice during development with those same populations that were increased in the ASD cohort. This is necessary, considering that ASD is a neurodevelopmental condition, and therefore developmental effects of microbiome need to be determined for their effects on ASD related behavior. In the current study, we analyzed the gut microbiome in individuals with ASD and controls, from an Israeli cohort of children and youth. This was followed by subsequent treatment of mice during development with a bacterial genus that was over-represented among the individuals with ASD, followed by behavioral and molecular analysis of the mice at eight weeks of age.

## Results

In order to determine possible differences in microbiome composition associated with ASD, we performed 16S rRNA amplicon sequencing on fecal microbiome samples from 42 non-sibling controls and 96 individuals diagnosed with ASD in Israel. There were no significant differences in age and sex between groups (average ages: control 7.4±SD3.2, Autism 8.7±4.1 p=0.082, Sex ratio: Control group 30 males 12 females, Autism group 72 males 25 females. fisher’s exact test p=0.83).

Comparing beta diversity (between samples) across groups, calculated using weighted unifrac distances, revealed a significant difference between the control and ASD samples (p=0.002 PERMANOVA) (Figures 1A, B). Alpha diversity (within samples) comparisons, using Faith’s phylogenetic distance (PD) demonstrated a significant increase in diversity in the ASD samples (p=0.005) (Figure 1C). These initial analyses support differences in the diversity of the microbiome in individuals with ASD.

**Figure 1.**
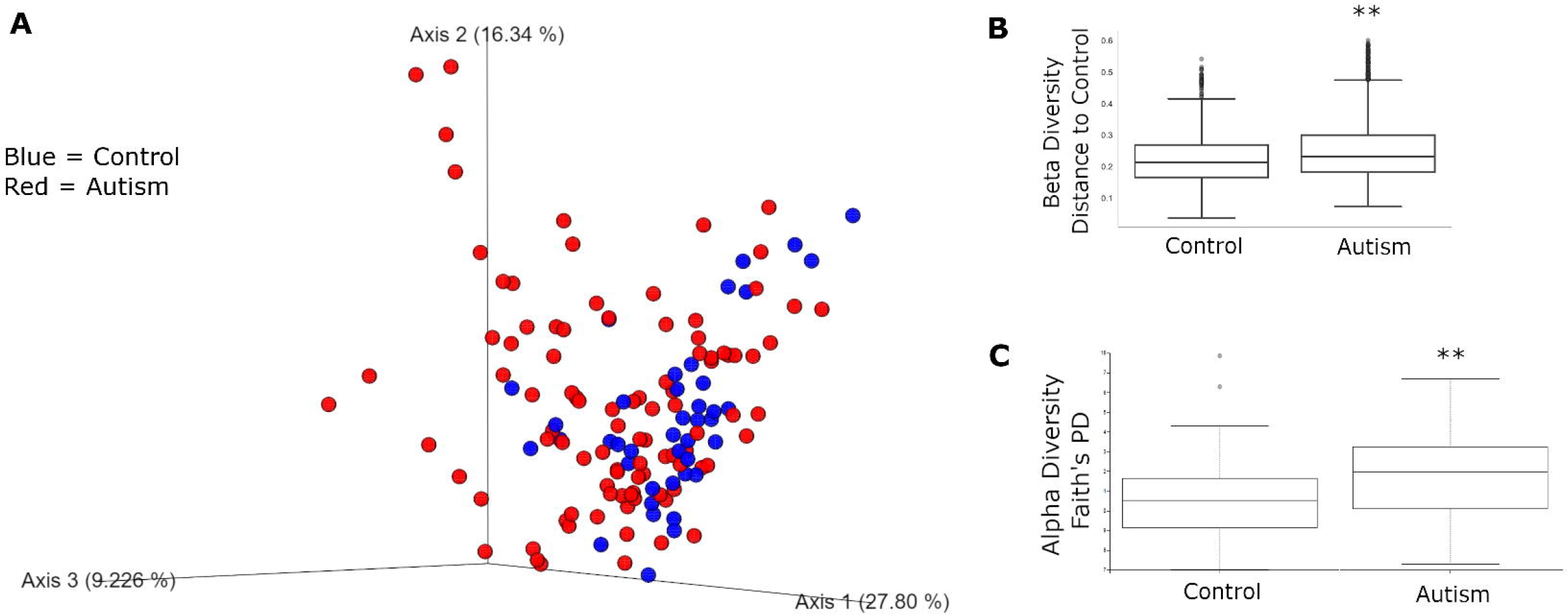
Beta and alpha diversity of microbiome compared according to diagnosis with ASD. A.) PCoA plot of all samples. Control samples are in blue while ASD samples are in red. B.) Beta Diversity Analysis as differences from each sample to control samples. There is a significant difference in beta diversity between ASD and control groups (PERMANOVA p=0.002). Whisker plots representing quartiles. C.) Alpha-Diversity analysis using Faith’s PD. Alpha Diversity is increased in ASD group (PERMANOVA p=0.005) Whisker plots representing quartiles. **=p<0.01

To determine the differences in taxa while considering both age and sex as covariates, we performed Aldex2 (Fernandes et al., 2014), which is a bioinformatic pipeline designed specifically for microbiome data. Defining age and sex as random effects, we built a general linear model^13^ . At a cutoff of FDR<0.1, The only significant differentially abundant phylum was Bacteroidetes (p=0.0052, FDR =0.048), while the only significant differentially abundant genus was *Bacteroides* (p=0.00093, FDR=0.095), a member of the phylum Bacteroidetes, both of which were relatively more abundant in the ASD group. Cohort data included eating habits, particularly if the individual had a gluten-free diet, dairy-free diet, and vitamin supplementation. Among the autism cohort, 11 had gluten-free diets, 16 had dairy free diets, and 20 took vitamin supplementation. In comparison, only one person had gluten-free diet, one person took vitamin supplementation, and none had diary free diets in the non-sibling control cohort. Therefore, we also performed the general linear model while including these factors as random effects. Inclusion of eating habits had no effects on the statistical significance of Bacteroidetes (p=0.0048, FDR=0.045) (Figure 2A) and *Bacteroides* (p=0.0087, FDR=0.089) (Figure 2B). Therefore, the changes in these bacteria are directly associated with autism diagnosis with no effect of diet. To visualize the role of *Bacteroides* as a source of variance in the overall microbiome signature, we replotted the Principal Coordinate Analysis (PCoA) graph with levels of *Bacteroides* as the identifier (Figure 2C, darker green is more *Bacteroides*).

**Figure 2.**
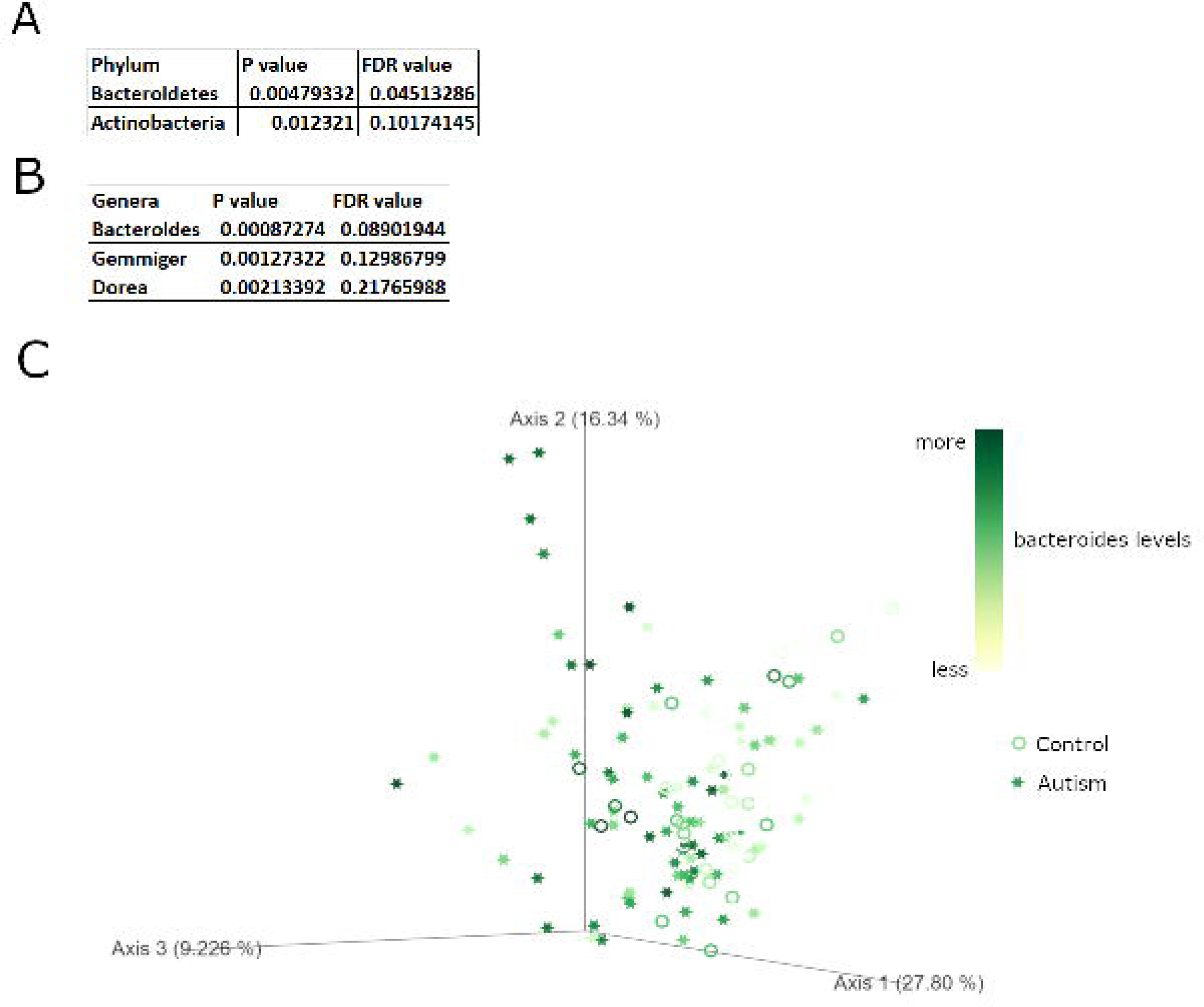
Differentially abundant microbial phylum and genera in ASD samples. A.) Differentially abundant bacterial phylum as calculated by general linear model with age, sex, and diet covariates. B.) Differentially abundant bacterial genera as calculated by general linear model with age, sex and diet covariates.C.) PCoA plot of all samples on a scale from white (low *Bacteroides* levels) to dark green (high *Bacteroides* levels). Autism samples in stars and control samples as circles.

The increase in *Bacteroides* in individuals with ASD was initially surprising, considering previous findings from mouse models found that one species of *Bacteroides, B. fragilis*, can attenuate repetitive behaviors in adult mice (Hsiao et al., 2013). However, we hypothesized that *B. fragilis* may have different effects on behavior and brain function during development. Therefore, to understand if presence of *B. fragilis* during a critical developmental window initiates ASD-related behaviors, we performed a one-time gavage of *B. fragilis* or vehicle (PBS) to newborn (P0.5) specific pathogen free (SPF) C57 mice, followed by determination of successful *B. fragilis* colonization in the gut and behavioral phenotyping in adulthood. Stools were collected at the age of two weeks and eight weeks for microbiome profiling. After two weeks, *B. fragilis*-treated animals displayed a different microbiome composition from control mice when examining beta diversity (Figure 3A). In addition, relative abundance of *B. fragilis* was significantly higher (by over a million-fold) in *B. fragilis* treated mice compared to control mice in both males and females (Figure 3B), as determined by quantitative PCR. However, these differences were not observed after eight weeks (Figure 3C, D). Therefore, *B. fragilis* treatment leads to a robust increase in bacterial levels by two weeks after treatment, but these levels decrease by adulthood.

**Figure 3.**
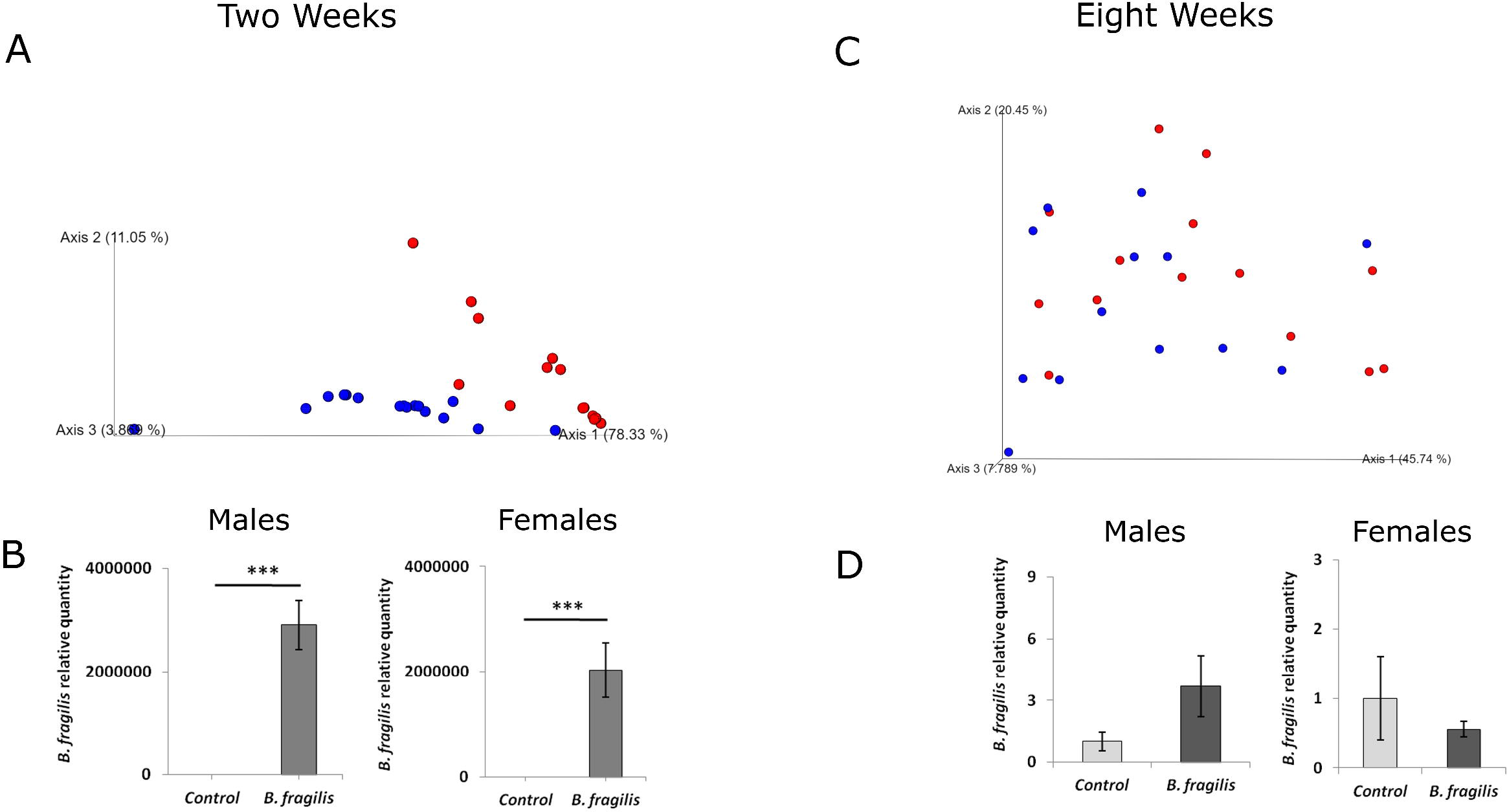
Examination of gut microbiome in mice after treatment of newborn pups with *B. fragilis*. Stool was taken from mice at ages of 2 weeks and 8 weeks and 16S rRNA amplicon sequencing for microbiome analysis was performed in parallel to quantitative PCR for *B. fragilis* abundance. A.) PCoA plot of weighted unifrac distances from microbiome at two weeks of age. Blue are non-sibling controls and Red are *B. fragilis* treated. B.) qPCR analysis of *B. fragilis* in stool of both males and females at two weeks of age (n=6, ***p<0.001, two tailed student’s t-test. Error bars represent standard error). C.) PCoA plot of weighted unifrac distances from microbiome from eight weeks of age. Blue are non-sibling controls and Red are *B. fragilis* treated. D.) qPCR analysis of *B. fragilis* in stool of both males and females at two weeks of age. n=6 Error bars represent standard error.

In order to determine if early administration of *B. fragilis* has a long-lasting effect on behavior in adulthood, behavioral tests were performed on eight-week-old *B. fragilis*-treated mice and control mice. Male *B. fragilis*-treated mice buried significantly more marbles in the marble burying test (Figure 4A). Considering that this finding may be indicative of increased repetitive behaviors, we performed an additional test of repetitive behaviors, calculating time spent grooming in an open field. Male *B. fragilis*-treated mice spent significantly more time grooming (Figure 4B) compared to control mice. Male mice showed no differences in time spent in the center of the open field (Figure 4C) or total distance travelled in an open field (Figure 4D), measures of anxiety. In females, no differences were observed between the two groups in all four tests (Figure 4E-H). Thus repetitive behaviors are mediated by *B. fragilis* in a sex-dependent manner.

**Figure 4.**
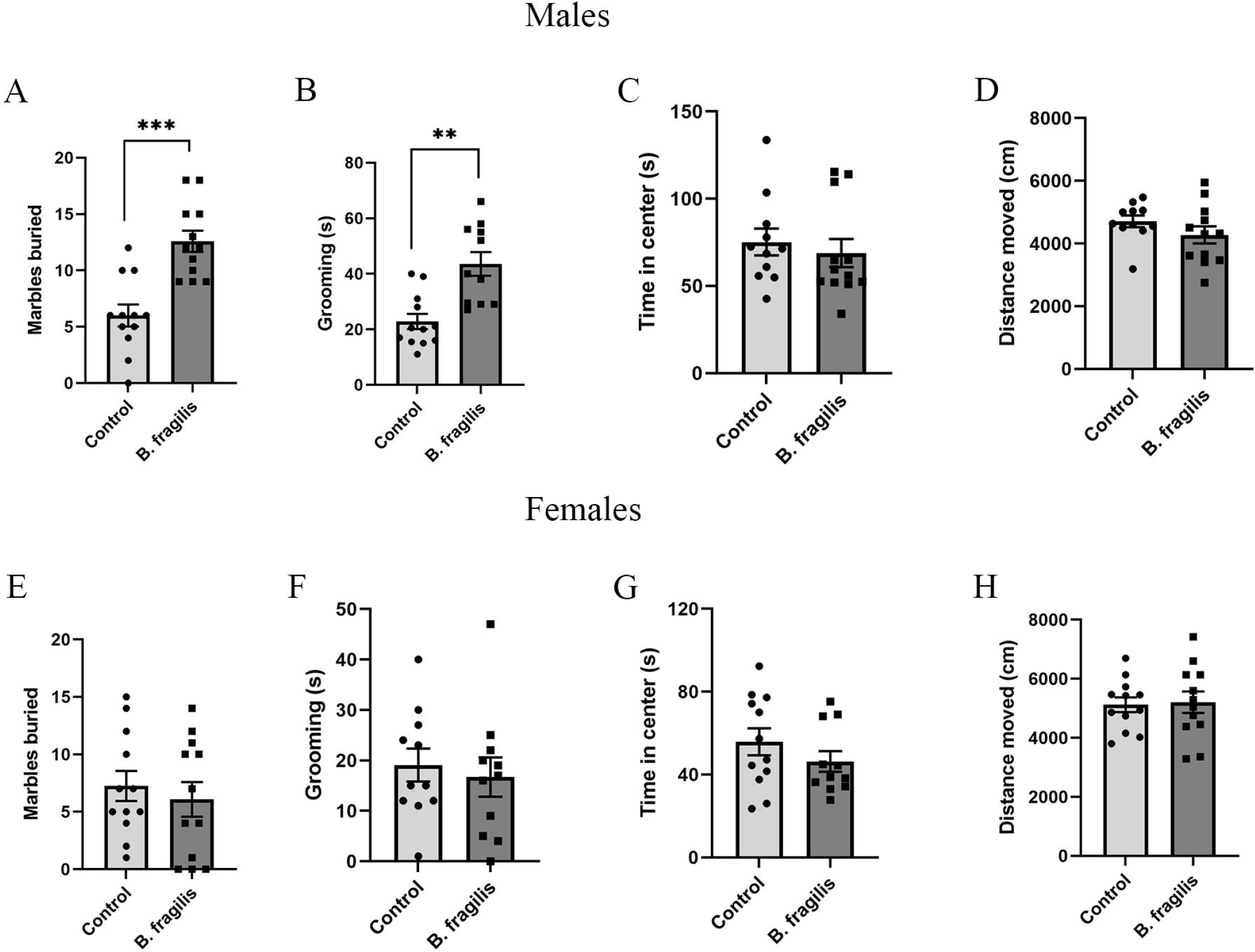
Dysregulation of adulthood repetitive behavior specifically in male mice treated with *B. fragilis*. Behavioral analysis of eight-week-old *B. fragilis*-treated mice. A.) Total number of marbles buried by male mice in a 30 minute period B.) Time spent grooming in an open field arena by male mice in a ten minute period in open field arena C.) Time spent in the middle of an open field arena by male mice in a 10 minute period. D.) Total Distance Traveled by male mice in an open field arena in a 10 minute period. E.) Total number of marbles buried by female mice in a 30 minute period F.) Time spent grooming in an open field arena by female mice in a ten minute period in open field arena G.) Time spent in the middle of an open field arena by female mice in a 10 minute period. H.) Total Distance Traveled by female mice in an open field arena in a 10 minute period. Two tailed student’s t-test **=p<0.01 ***=p<0.001 n=11 per group. Error bars represent standard error.

We next tested the effects of *B. fragilis* administration on performance in the three chambered social tests. In the sociability test, untreated and treated animals displayed the same preference to sniffing a novel mouse compared to an empty chamber. (Figures 5A, C). However, in the social novelty test *B. fragilis*-treated males showed less preference towards interacting with a novel animal compared to untreated males (Figure 5B). In females, though, both the treated and untreated groups displayed normal sociability and social novelty behavior (Figure 5C-D). In conclusion, males treated with *B. fragilis* demonstrated more repetitive behavior and less preference towards social novelty in comparison to untreated males.

**Figure 5.**
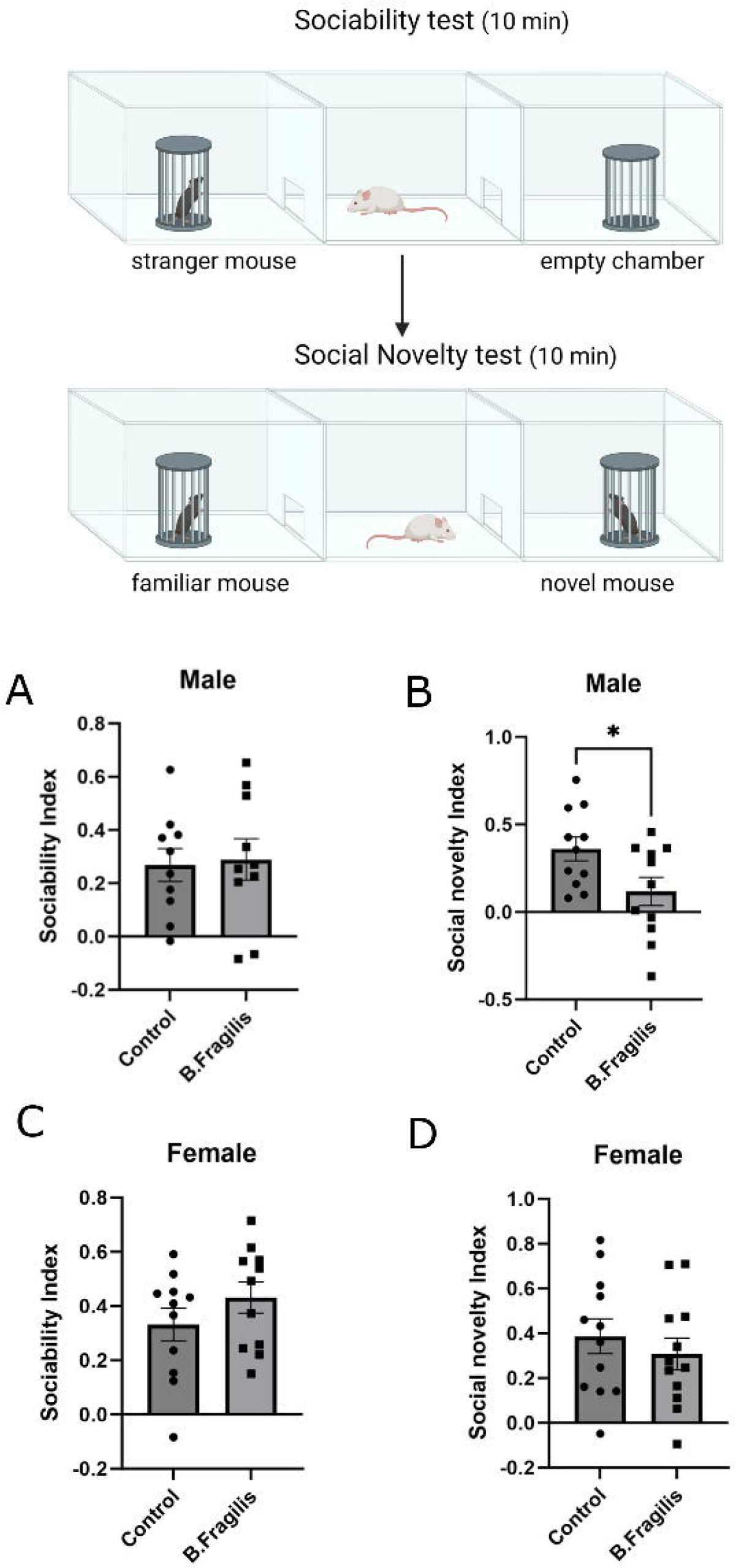
Male mice treated with *B. fragilis* display dysregulated social novelty behavior. Behavioral analysis of eight-week-old *B. fragilis*-treated mice in three chamber social behavior paradigm. A.) Preference of male mice for sniffing an empty chamber or chamber with a novel mouse. Calculated as social index. B.) Preference of male mice for sniffing a chamber with familiar mouse or novel mouse. Calculated as social novelty index. C.) Preference of female mice for sniffing an empty chamber or chamber with novel mouse. Calculated as social index. D.) Preference of female mice for sniffing chamber with familiar mouse or novel mouse. Calculated as social novelty index. Two tailed paired student’s t-test **=p<0.01 ***=p<0.001 n=11 per group. Error bars represent standard error.

Considering that behavioral changes were present in adult offspring, we next tested if the newborn *B. fragilis* treatment affected cortical gene transcription in adulthood. The prefrontal cortex was dissected from male adult mice, and global gene expression analysis was performed by RNA-seq. No genes passed differential expression testing at a threshold of FDR<0.1 (Figure 6A). We further explored the data using Gene Set Enrichment Analysis (GSEA), which can uncover gene biological pathways that are associated with the more upregulated or downregulated genes within a gene expression dataset^14^. Genes were ranked by their fold change from most increased in *B. fragilis* treated group to the most decreased genes. We found a significant enrichment of increase cholesterol biosynthesis related genes among genes that are increased in the *B. fragilis* group (Figure 6B) and downregulation of genes regulating neural crest differentiation (Figure 6C). When comparing our data to databases of genes found in specific cell types, we found a significant downregulation of genes expressed in inhibitory interneurons (Figure 6D) and inhibitory neuron progenitors (Figure 6E). These findings suggest that overabundance of *B. fragilis* in key development stages (here, in newborn pups) can have a long-term impact on cortical gene expression at the network level, but not at the level of specific genes

**Figure 6.**
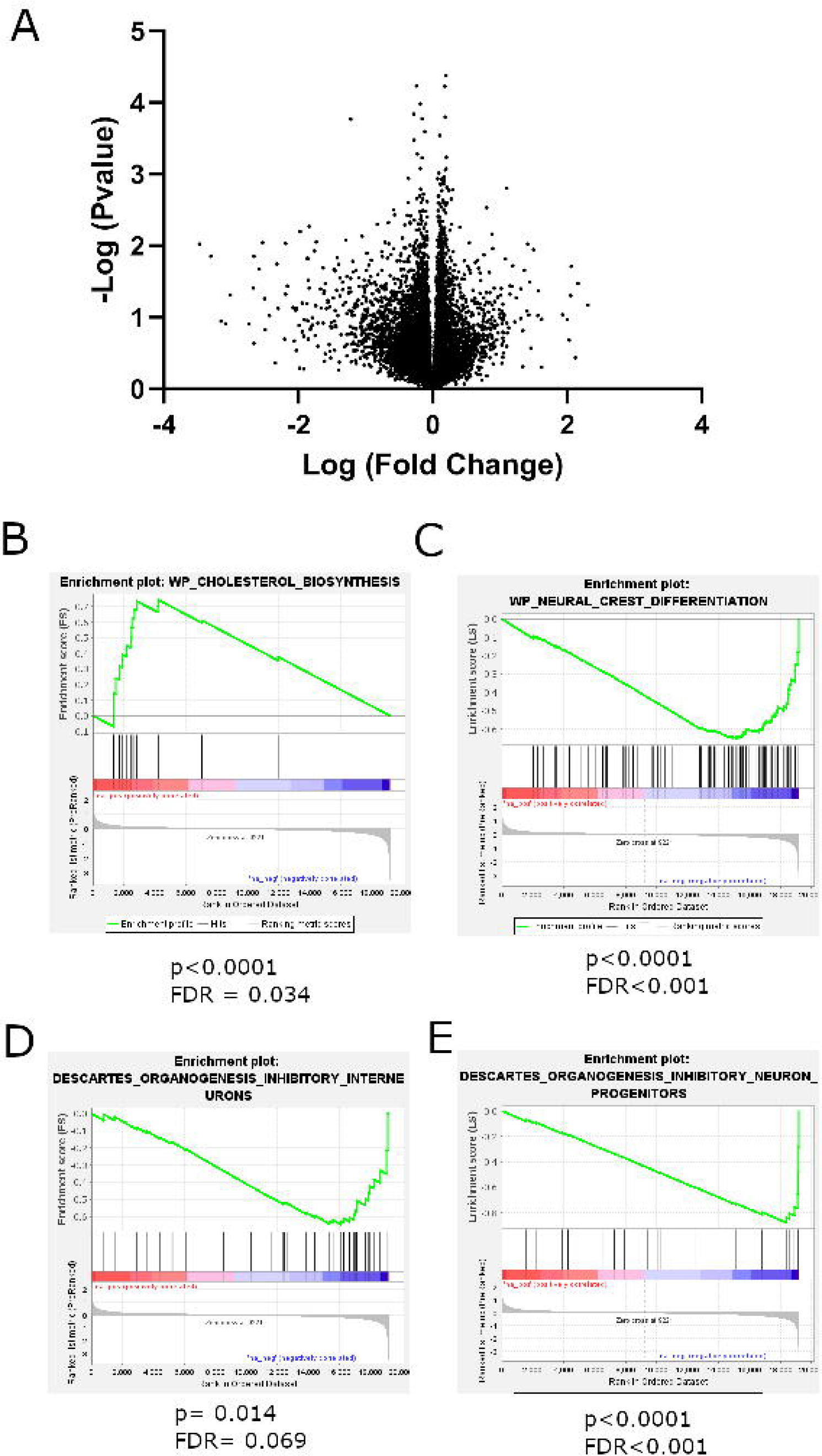
Whole transcriptome analysis of adult male prefrontal cortices in *B. fragilis* treated mice. A.) Volcano plot of genes in differential expression analysis between *B. fragilis* treated and untreated mice. X axis is log fold change of genes between treatment groups and Y axis is negative log p value of each gene. B-E.) GSEA analysis describing enriched gene categories among upregulated or downregulated genes. B) Enrichment of cholesterol biosynthesis genes in *B. fragilis*-treated group and C) a decrease of genes related to neural crest differentiation in the same treatment group. D) A decrease in genes expressed specifically in inhibitory interneurons in the *B. fragilis*-treated group together with a decrease in E) inhibitory interneuron progenitor genes in the same group.

## Discussion

In this study, we identified changes in the microbiome in individuals diagnosed with ASD in an Israeli cohort. The main differences were an increase in alpha diversity and an increase in the relative abundance of the phylum Bacteroidetes and the genus *Bacteroides*. Both findings were surprising, although for separate reasons. Decreased alpha diversity has been associated with decreased health in several conditions. There is however some evidence that this is not the case in neurological diseases^15^. The increased alpha diversity seen in our ASD cohort is therefore somewhat surprising. In a recent study, we found that increased alpha-diversity in the microbiome of an ASD mouse model was associated with increased bacterial load^16^. Therefore, we cannot rule out the possibility that these individuals have an increased bacterial load. The increase in *Bacteroides* was initially rather surprising considering a previous rodent study demonstrating that a species of *Bacteroides, B. fragilis*, may attenuate ASD-associated behavior^10^. In that study, maternal immune activation in mice induced ASD-like behavioral abnormalities in pups, in association with decreases in *B. fragilis* abundance. Treatment of adult offspring with *B. fragilis* attenuated the ASD-like behavioral phenotypes. However, recent studies have also determined increases in *Bacteroides* in individuals with ASD. In a meta-analysis using several cohorts, Morton *et. al*. found an increase of *Bacteroides* in ASD, that also correlated with increases in interleukin-6^17^. The effects of this species could be solely exposure time dependent (infancy vs adulthood), as demonstrated here, or they could also be strain specific.

In our experiment, *B. fragilis* administered in newborn pups was able to colonize the gut for at least two weeks after treatment but there were no significant changes between experimental groups in adulthood. Despite its absence in adulthood, behavioral effects were observed in a sex-specific manner (males only). When looking at gene expression in adult male prefrontal cortices, we found downregulation of genes specifically found in inhibitory interneurons. Dysregulation of the excitatory/inhibitory balance in the cortex, and specifically an increase in excitatory signaling, has previously been indicated as a possible biological driver of ASD^18,19^. In addition, there is an increase in genes associated with cholesterol biosynthesis. Previous studies have found dysregulation of serum cholesterol homeostasis in individuals with ASD, and hypocholesterolemia is the most common finding^20^. However, there is still little knowledge in regard to brain cholesterol homeostasis in ASD. A recent study found that the FMR1 mouse model of autism displays differences in cholesterol receptor expression in the brain, with Low-density Lipoprotein Receptor expression increased specifically in the cortex^21^. This correlates well with our finding of increased cholesterol biosynthesis genes in the cortices of *B. fragilis* treated mice. Dysregulation of cholesterol, as a crucial regulator of membrane biology and signal transduction, may affect the proper function of neuronal signaling in these autism models. Therefore, these findings point to a possible biological mechanism through which *B. fragilis* induces ASD-like behavior. However, it is still not clear how these changes in gene expression take place.

While it is not clear why B. fragilis did not maintain their high levels in gavaged mice until adulthood, it is possible that this is due to the increasing microbial diversity during development. Previous studies in humans found that newborns have a relatively low microbial diversity, which increases linearly with age^22,23^. Therefore, it is possible that B. fragilis was able to colonize the gut microbiome during early developmental time points, but was not able to compete with other microbial species as the microbiome diversified. Additional support for this theory is a separate study that found that babies who had an increase in *Bacteroides* in the first week of life, due to cesarean section, displayed normal levels of *Bacteroides* by the second week of life^24^ (PMID: 33377127). While these findings are in a different species and time scale, this further suggests that artificially increased levels of *Bacteroides* will eventually return to normal levels.

While members of the genus *Bacteroides*, including *B. fragilis*, have many positive roles in human health, there are also possible downsides of overabundance of these species^25^. Specific strains of *B. fragilis* excrete a toxin known as *B. fragilis* toxin (BFT), which is a metalloprotease capable of cleaving E-cadherins^26^. As such, this toxin increases gut permeability and may lead to leaky gut. There is additional evidence that while *Bacteroides* is usually a symbiotic genus, it may cause infections under specific circumstances^27,28^. Therefore, further work is necessary to understand if *B. fragilis* can induce specific changes in permeability or other gut parameters in the context of ASD.

The sex-specificity of the behavioral and molecular findings in mice are of great interest, considering the male bias of ASD diagnosis. There is much debate regarding the source of the male bias in autism diagnosis, and our findings further indicate that males may have a particular susceptibility to environmental causes of ASD. Of further interest, it has been found that males have a higher abundance of Bacteroides than females^29^. Therefore, it is possible that males are more susceptible to the effects of a further increase in Bacteroides abundance.

A previous study demonstrated that maternal immune activation during pregnancy in mice induces ASD-like behaviors in offspring, in addition to decreases in levels of *B. fragilis*^10^. Furthermore, they demonstrated that ASD-like behaviors could be attenuated by treatment with *B. fragilis* in adulthood. In our current study, we show that one-time treatment of newborn mice with *B. fragilis* induces ASD-like behaviors. The contrasting findings between our studies are likely due to different effects of *B. fragilis* in development versus adulthood. In our study, *B. fragilis* colonized the mice for up to two weeks after treatment but was no longer present at eight weeks; however, behavioral and molecular phenotypes remained, suggesting that *B. fragilis* can affect proper brain development. The current study adds to recent evidence that increase of *Bacteroides* is found in cohorts of individuals with ASD and that it may play a role in regulation of neurodevelopment.

## Acknowledgements

We would like to thank Teva Pharmaceuticals for sponsorship of the study as part of its support for the Bar Ilan University Faculty of Medicine. EE is further supported by a grant from the Israeli Science Foundation (1159/22). OK is supported by the European Research Council (ERC) under the European Union’s Horizon 2020 research and innovation programme (Grant agreement ERC-2020-COG No. 101001355).

## Materials and Methods

### Human Cohort

All individuals were recruited to this study through the Israel Autism Biobank and Registry in the Azrieli Faculty of Medicine at Bar Ilan University. This biobank and registry are approved through the Israeli Ministry of Health, and this study was approved with the Helsinki approval of Ziv Hospital in Safed (protocol 0090-2018-ZIV). All participants signed letter of consent for this protocol.

The ASD cohort includes 96 individuals diagnosed with ASD. Exclusion criteria included other mental diagnoses. All stool samples were taken at least three months after last use of antibiotics. Stool samples were collected with a home stool collection kit given to families. After collection, a delivery company brought the samples to the Azrieli Faculty of Medicine at Bar Ilan University (Safed, Israel), and samples were immediately stored at -80°C. No more than two hours passed between sample collection and -80°C storage.

### Microbiome sequencing and pre-processing

DNA was extracted from stool samples using the PureLink™ Microbiome DNA Purification Kit (ThermoFisher, Waltham, MA, USA) according to the manufacturer’s instructions and following a 2-minute bead beating step (BioSpec, Bartlesville, OK, USA). Purified DNA was used for PCR amplification of the variable V4 region using the 515F and 806R barcoded primers following the Earth Microbiome Project protocol (Human Microbiome Project Consortium, 2012). For each PCR reaction, the following reagents were added: 4μl (20ng/ μl) DNA (sample), 2 μl 515F (forward, 10μM) primer, 2 μl 806R (reverse, 10μM) primer, and 25 μl PrimeSTAR Max PCR Readymix (Takara, Mountain View, CA, USA). PCR reactions were carried out by 30 cycles of denaturation at 98°C for 10 seconds, annealing at 55°C for 5 seconds, extension at 72°C for 20 seconds and then a final elongation at 72°C for 1 minute. Amplicons were purified using AMPure magnetic beads (Beckman Coulter, Brea, CA, USA) and further quantified using the Picogreen dsDNA quantitation kit (Invitrogen, Carlsbad, CA, USA). Then, equimolar amounts of DNA were pooled and sequenced using the Illumina MiSeq platform and MiSeq Reagent Kit V2 (500 cycles) at the Bar-Ilan University, Azrieli Faculty of Medicine Genomic Center.

Microbial diversity and composition were assessed using QIIME2 version 2019.4^30^. First, single-end sequences were imported (qiime import) and demultiplexed (qiime demux) with golay error correction. Next, sequences were denoised using DADA2^31^ (qiime dada2 denoise-single), trimming the first 5 bases and truncating each sequence at position 215. A phylogenetic tree was constructed using the fragment-insertion method (qiime fragment-insertion sepp). Taxonomic classification was done using a naïve-based classifier trained on the 99% Greengenes 13_8 V4 reference set (qiime feature-classifier classify-sklearn). In order to remove low-confidence features, only features with a frequency higher than 50 reads in at least 5 samples were kept. Differences in relative abundances of bacterial taxa between groups were identified using the ANOVA-like differential expression tool (ALDEx2)^13^. Sex, age, and dietary patterns were defined as covariates in this analysis of the human cohort.

### Mouse Housing

C57BL/6J (C57) breeder mice purchased from ENVIGO (Rehovot, Israel) were used in this study. The mice were kept in SPF conditions, housed under reverse 12h light/dark cycle, provided with water and food. Mice were weaned 4 weeks after their births and group housed with 3-5 animals per cage. There are at least three cages per experimental group. Animals that developed illnesses during the experiments were excluded. All procedures were approved by the Animal Ethics Committee of the Bar-Ilan University, Israel (protocol number 36-06-2019).

### Bacterial treatment

*Bacteroides fragilis* (*B. fragilis*) obtained from DSMZ (*Bacteroides fragilis* DSM 2151) was used in this study. *B. fragilis* was cultivated in Cooked Meat Broth with hemin and vitamin K in anaerobic conditions for 24h (optical density of ∼0.5 at 600nm). Media and reagents were obtained from SIGMA-ALDRICH (Merck Darmstadt, Germany) and prepared according to manufacturer’s instructions. On the day of treatment, all the bacteria were washed three times with PBS (by centrifugation, taking out supernatant, adding PBS, pipetting and vortexing). Pregnant mice were checked for deliveries each day, and during the first day after birth, *B. fragilis* or PBS was administered to the pups by gavage. Gavage was given under a heating lamp via a 2.5cm segments of polyethylene tubing PE10 (.011x.024 inch) attached to disposable insulin syringes. For each mouse, 10^9^ bacterial colony-forming units (CFU) were suspended in 10μl of sterile PBS. Control groups were given the same volume of sterile PBS.

### Feces collection and DNA extraction for mice

Mice fecal samples were collected on week 2 and 8 (before behavioral assays) and stored in -20ºC or -80ºC. Microbial DNA was isolated from the stool samples as described above. DNA was quantified using a Qubit dsDNA HS assay kit (Life Technologies, Carlsbad, CA, USA) according to the manufacturer’s protocol and as described previously^32^.

### Real-time quantitative PCR (qRT-PCR)

Abundance *of B. fragilis* in stool was determined by ViiA™7 Real-Time PCR System (Life Technologies), in 10 μl volumes. qRT-PCR consisted of 40 cycles, melting temperature of 95°C for ten seconds per cycle, and annealing/elongation temperature of 60□°C for thirty seconds per cycle. It was performed using Fast Start Universal SYBR Green Master (Rox) (Roche, Basel, Switzerland) and primers for 16S *a*nd *B. fragilis* (primer sequences are indicated in Table 1) at a final concentration of 250nM each. Total 16S DNA was used as an endogenous control. All samples were tested in triplicates. In addition, *B. fragilis* was detected by visualizing PCR bands after running DNA samples that were amplified by qPCR on a 2% agarose gel.

### Behavioral tests

Behavioral tests were conducted at age of 9-12 weeks. All tests were done during the dark phase, and mice were acclimatized to the behavior room for at least 1 h before testing. Each test was performed on a separate day, usually with one day rest between each test. Experiments were recorded with the Panasonic WV-CL930 camera and with the Ganz IR 50/50 infrared panel. The recorded movement of the mice was analyzed by the Ethovision XT 10/11 (Noldus, Wageningen, Netherlands) software.

### Open Field test

The open field test examines anxiety-like behavior in mice, by testing the conflict between the desire to explore a new open space (by entering the center of the box) and the desire to stay protected (staying in the corners). It also examines locomotion by the total distance moved. The mouse is placed in the corner of a plastic square box (50x50) under light of 120 lux where it moves freely for 10 minutes in order to examine anxiety and locomotion. An extra 10 minutes are added in order to assay grooming. The time of self-grooming is calculated for the second 10-minute block in order to examine repetitive-like behavior.

### Marble burying test

The marble burying test examines repetitive-like behavior. In a non-glare Perspex arena (20X40 cm), twenty green glass marbles (15 mm in diameter) are arranged in a 4X5 grid that covers 2/3 of the apparatus on top of 5 cm of clean bedding. Each mouse is placed individually in the corner that does not contain marbles and is given a 30-minute exploration period, after which the number of buried marbles is counted. “Buried” is defined as 2/3 covered by bedding. Testing is performed under dim light (25 lux).

### Three-chamber social interaction test

In the three-chamber social interaction test, the test mouse is placed in the middle chamber of a three-chamber apparatus. In each side chamber, there is a small container that can be left empty or filled with a mouse. The social interaction test measures sociability by examining the preference of the mouse to sniff a novel mouse within a container or to sniff an empty container. The social novelty test measures novelty by examining the preference between sniffing the first stranger (familiar mouse) or the new stranger (novel mouse). The mouse is placed in the middle chamber for habituation (5 min), during which entry to both side chambers is barred. After habituation, the mouse is given 10 minutes to move between chambers, and sociability is scored using the Ethovision software (Noldus, Wageningen, Netherlands). Social index is calculated as (time sniffing social chamber -time sniffing empty chamber)/(time sniffing social chamber+time sniffing empty chamber). Social novelty index is calculated as (time sniffing novel mouse chamber -time sniffing familiar mouse chamber)/ (time sniffing novel mouse chamber + time sniffing familiar mouse chamber).

### Brain dissection and mRNA sequencing

Ten week old male mice were sacrificed by rapid decapitation and brains were quickly removed. Punches were taken from prefrontal cortex and kept in centrifuge tubes at -80°C. RNA was extracted from brain regions with a Qiagen RNAeasy mini kit (Qiagen, Hilden, Germany). There were eight mice in each experimental group. Total RNA was extracted using the Qiagen miniRNAeasy kit (Qiagen, Hilden, Germany). From 100□ng of total RNA of each sample, mRNA enrichment was done with the NEBNext Poly(A) mRNA magnetic isolation Module (NEB # E7490) (New England Biolabs (NEB), Ipswich, MA, USA), followed by preparation of RNA libraries using the NEBNext Ultra RNA Prep kit (NEB #E7530) (New England Biolabs (NEB), Ipswich, MA, USA), according to manufacturer’s protocols. Libraries’ concentrations were determined by Qubit (ThermoFisher, Waltham, MA, USA), while quality and size distribution were analyzed using Bioanalyzer 2100. The sequencing was performed at the Bar-Ilan University, Azrieli Faculty of Medicine Genomic Center. The quality of the sequenced data, as well as read length distributions after trimming, was evaluated by FASTQCT (0.11.5). Bioinformatic analysis was performed at the Mantoux Bioinformatics Institute of the Nancy and Stephen Grand Israel National Center for Personalized Medicine at the Weizmann Institute of Science. The reads were mapped to the *Mus musculus* reference genome, GRCm38.p5, using the Tophat2 software. An average of 20 million reads were obtained per sample. Deseq2 software was used for differential expression analysis. GSEA software was used for gene network analysis^14^. Genes were preranked by logfold2 change and then subject to GSEA analysis.

### Statistics

In animal experiments, all statistics were performed in Graphpad prism (San Diego, CA, USA), using the appropriate statistical tests after testing for equality of variances. Other statistical tests, as appropriate, are described within the materials and methods section.

## Author Contribution Statement

J.C, N.G., and R.M. did sample preparation and experimentation on human samples. R.M. and D.G. performed mouse studies. J.C. and N.S. directly managed the project. E.T. performed bioinformatic analysis. M.S., S.K., A.K., N.K., T.K., and T.B. were involved in sample procurement and management. O.K. and E.E. were involved in study design. E.E. and J.C. wrote the manuscript. All authors reviewed the manuscript.

